# Effects of auditory noise intensity and color on the dynamics of upright stance

**DOI:** 10.1101/2023.12.12.571330

**Authors:** Sam Carey, Jessica M. Ross, Drew Abney, Ramesh Balasubramaniam

## Abstract

Previous work assessing the effect of additive noise on the postural control system has found a positive effect of white noise on postural dynamics. This study covers two separate experiments that were run sequentially to better understand how the structure of the additive noise signal affects postural dynamics, while also furthering our knowledge of how the intensity of auditory stimulation of noise may elicit this phenomenon. Across the two experiments, we introduced three auditory noise stimulations of varying structure (white, pink, and brown noise). Experiment 1 presented the stimuli at 35 dB while Experiment 2 was presented at 75 dB. Our findings demonstrate a decrease in variability of the postural control system regardless of the structure of the noise signal presented, but only for high intensity auditory stimulation.

## Introduction

Postural control has been the subject of scientific interest for many years due to the complexity of the human postural system and its interactions with the environment. Changes in environmental contexts such as: changes in cognitive load (Balasubramaniam and Wing, 2002), external sensory input (Ross et al., 2015, 2016; Priplata et al., 2002, 2003, and 2006), or the addition of secondary motor movements (Balasubramaniam and Wing, 2002) can alter the dynamics of balance. Despite the challenges presented by navigating through an ever-changing environment, the human postural system is capable of adapting to environmental variability quite efficiently. However, the specific processes by which external stimuli are filtered or processed during upright standing are not yet fully understood.

Postural control is a perceptual motor process that utilizes a continuous stream of sensory input from the auditory, somatosensory, vestibular, and visual systems to maintain a stable body position during standing (Dozza et al., 2007; Hegeman et al., 2005; Palm et al., 2009). Human balance relies on this redundancy of sensory input to account for possible changes or perturbations in the expected sensory information from these systems. This process requires the processing of both internal and external sensory information to detect any perturbations that may threaten balance and select the necessary motor responses needed to maintain stability (Schmidt, 1975). For instance, while the eyes are open, humans rely primarily on visual feedback for balance, but when visual input becomes limited or hindered, we rely more on somatosensory stimulation, such as through a light touch to the finger, to maintain an upright position (Wing, 2011). Furthermore, studies have demonstrated that additional sensory input through the auditory (Dozza et al., 2007), somatosensory (Priplata et al., 2003), vestibular (Hegeman et al., 2005), or visual modalities (Palm et al., 2009) can beneficially influence the maintenance of postural stability.

Understanding the sensitivity of the postural system and the ways in which external and internal information influences postural dynamics holds promise for individuals who are at an increased risk of falls. Past work has shown how increases in external sensory information can decrease sway variability and increase stability (Collins et al., 1996; Jeka et al., 1997; Priplata et al., 2003; Deviterne et al., 2005; Dozza et al., 2007; Palm et al., 2009; Ross et al., 2015, 2016). However, much remains unknown regarding not only how the presence or absence of new sensory information may influence motor dynamics, but also how the intensity or type of information present may differentially influence these systems. One potential theory to explain why additive sensory information may alter postural dynamics is Stochastic Resonance.

Stochastic Resonance (SR) is a phenomenon observed in nonlinear systems when the addition of noise to a system results in an optimal level of information transfer and an increase in output performance (Benzi, 1981, Hänggi 2002). Originally developed for modeling the cycle of glaciation on the earth, SR has since been observed in natural and experimental settings. The theory assumes that within a threshold-based system, any underlying information carrying signals within that system can become enhanced through the addition of noise onto the original signal. The addition of noise is assumed to increase the amplitude of the underlying signal, allowing for an increase in the frequency of threshold crossings necessary to send the information the signal is carrying.

SR has been studied in humans in the context of sensory processing (Hidaka, Nozaki and Yamamoto 2000), including in auditory (Morse and Evans, 1996; Mangiore, 2012), visual (Simonotto, Riani, Seife, Roberts, Twitty, and Moss, 1997) and tactile perception (Collins, Imhoff, and Grigg, 1996; Richardson, Imhoff, Grigg, and Collins, 1998). In the field of postural control, SR has been used to investigate the impact of noise on sensory information processing and postural stability, with studies showing that the addition of noise can improve postural control in healthy individuals (Ross et al., 2015; Priplata et al., 2002, 2003, and 2006) and aging populations (Ross, et al., 2016). Thus, the application of SR to the study of postural control provides a potential avenue for developing interventions to improve balance and reduce the risk of falls in high-risk populations.

One of the major properties of SR is the optimization curve of the intensity of the additive noise. An optimal amount of noise results in the maximal enhancement of behavioral performance, whereas further increases in intensity can degrade the output performance of the system of interest (Moss, Ward, and Sannita, 2004). Similarly, too little noise can add no benefit of information transfer (cite?). The intensity of noise may play a more critical role than originally thought when the modality of noise input is considered. Ward et al., (2001) distinguished the optimal level of intensity of stimulation during additive noise of the visual, auditory, and tactile modalities. Similarly, during tactile stimulation, Priplata and colleagues (2003) were able to elicit a beneficial response to postural sway with sub-threshold stimulation at the bottom of the foot. Although a major assumption of SR work is that the noise added to the system of interest refers to ‘white noise’, there many different types of noise, or more specifically, degrees of noise that are perceivable to the human sensory system. This understanding led us to consider the usefulness of the varying degrees of noise to our sensory system and assess if the structure of the noise signal would affect our motor system differentially.

In signal processing, white noise is a random signal that has equal intensity across all frequencies. This means that every frequency within the audible range has the same power, resulting in a sound that is uniformly loud. Pink noise, on the other hand, has a spectral power density that decreases as frequency increases. As frequency increases, the amplitude of the sound decreases, resulting in a sound that has more bass and less treble. Brownian noise, or commonly known as brown noise, as we will refer to it in this paper, has a type of ‘random walk’ in which the value of the signal at any given point is the sum of the previous value and a normally distributed random value. Brown noise is a type of noise that is considered random, but its value at any given point is dependent on the previous value, resulting in a sound that is more complex and changes over time. The potential differences between white, pink, and brown noise may be influential to postural control because they represent distinctive frequencies and distributions of noise signals that may affect the postural control system differently. For example, white noise has equal energy per frequency interval, while pink noise has more power in lower frequency ranges. Brownian noise, on the other hand, has a power spectral density that decreases with frequency, and these low-frequency disturbances may have more significant effects on the postural control system than white noise. Understanding the effects of different types of noise on postural control can assist future work to investigate the postural control system’s sensitivity to the structure of sensory inputs and disturbances. It can also provide insight into how different sensory inputs may be used to maintain balance and how the postural control system adapts to changes in the environment.

In the current study, we examined sway variability during four auditory conditions: silence, white noise, pink noise, and brown noise. All noise conditions were presented with and without visual input. We presented these stimuli across two different experiments, one at a low intensity (35 dB) and one at a high intensity (75 dB). We did this to begin to uncover if the strength and structure of the additive noise are crucial to elicit a reduction in balance variability that has been shown in previous studies (Ross et al., 2015, Ross et al., 2016; Carey, Ross, and Balasubramaniam 2023). We hypothesized that different intensities of noise would have the same variability reducing effects on postural sway as seen in past work that used sub-threshold tactile stimulation as the locus of stimulation input (Priplata et al., 2002, 2003, 2006). Similarly, we expected there to be a reduction in postural sway variability while listening to all three different noise stimuli (white, pink, and brown), when compared to silence. However, we expected the structure of the noise signal to have an impact on the postural dynamics during stimulation. Previous research has shown the influence white noise has on postural sway, but not on how differently structured noise signals may impact sway dynamics. Due to the structure of brown noise having a type of random walk pattern similar to the movement of the center of pressure during upright standing (Collins and DeLuca 1993;1994), we expected a reduction in sway variability to be greatest during brown noise stimulation compared to the other noise conditions. Similarly, we expected pink noise to have the same magnitude of effect, or a more beneficial effect compared to white noise due to the lack of structure of white noise and pink noise being closer to brown noise which has similar dynamics to postural sway.

Past work by Carey et al. 2023 has shown the ability to induce this effect through the auditory and tactile modalities to similar degrees. To further understand the phenomenon observed in previous work on noise and the motor system, this current study introduced three noise signals of varying structures (white, pink, and brown) to the motor system to better comprehend how the noise signal is influencing motor behavior. We hypothesized that all three noise signals would have a positive effect on postural sway dynamics per the Stochastic Resonance phenomenon. By introducing the varying structures of noise, we hoped to be able to understand if frequency matching occurs between the noise signal and our postural control system based on the structure of the signal. If brown noise has the largest effect on balance variability, this suggests that there may be an mechanism at play other than SR. If all three noise signals have the same magnitude of effect on balance, this would support that frequency matching may not have a functional impact on balance variability and that sway reducing effects of noise on balance are better explained with SR or shifting auditory attention.

### Experiment 1

Experiment 1 attempts to extend pervious work on the influence of auditory noise signals on postural sway dynamics by adding noise with varying frequency content compared to previous work that utilized white noise only. This study aimed to replicate the additive noise effect but at a lower intensity than past work (35 dB) and extend to additional types of noise (brown and pink in addition to white). The goal of the study was to establish the effects of differently structured noise signals effects on postural sway when presented at a low sound intensity.

## Results

### Radial Sway

Radial sway was reduced with vision (Fig. 1). We observed a main effect of vision (F(1,21) = 26.36, η = .56, p = .001) and a main effect of condition (F(1,68) = 3.92, η = .16, p = .0125) on radial sway (RS; Fig. 1A and 1B). We did not observe any vision by noise interactions (F(1,21) = 1.12, η = .05, p = .347).

**Fig. 1.**
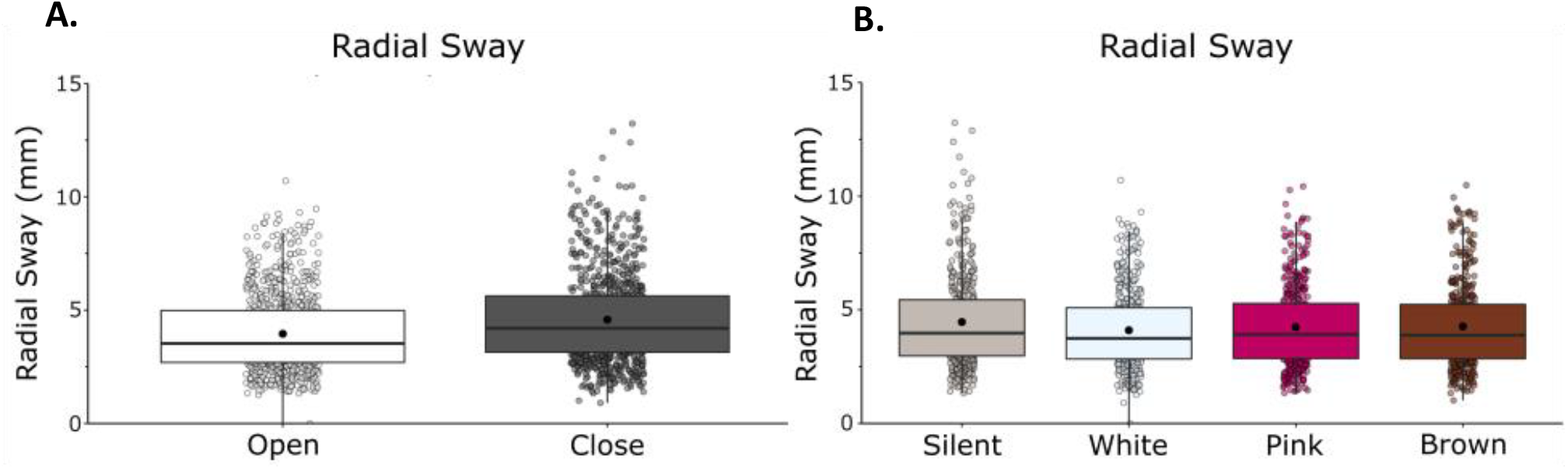
RS was significantly reduced with eyes open, but was unaffected by white noise, pink noise, or brown noise. A) RS in eyes closed/eyes open. B) RS in silent, white, pink, and brown noise conditions. There was no interaction between vision and condition. Box and whiskers plot with the solid black line representing the median, the solid black dot representing the mean, and the extending lines showing the maximum and minimum values.

Bonferroni-corrected post-hoc comparisons were performed to compare the individual noise condition effects on RS when compared to silence and to other noise conditions. Post-hoc comparisons revealed no significant difference between silence (M = 4.46, SD = 2.03) and white noise (M = 4.10, SD = 1.76, p = .096), pink noise (M = 4.24, SD = 1.78, p = .109) or brown noise (M = 4.27, SD = 1.83, p = .568). There was no difference between the stimulation conditions when compared to each other: white and pink (p = .873), white and brown (p = .708), pink and brown (p = 1.00). The significant main effect of condition is not supported by the post-hoc tests.

### High-Frequency Radial Sway

High-frequency RS was reduced with vision and noise (Fig. 2). We observed a main effect of vision (F(1,21) = 137.76, η = .87, p = .001) and a main effect of condition (F(1,21) = 2.91, η = .12, p = .041) on high-frequency RS (Fig. 2). We observed a vision by noise interaction effect, (F(1,21) = 4.38, η = .17, p = 0.007), which suggests that visual and auditory feedback contributed interactively to high-frequency sway.

**Fig. 2.**
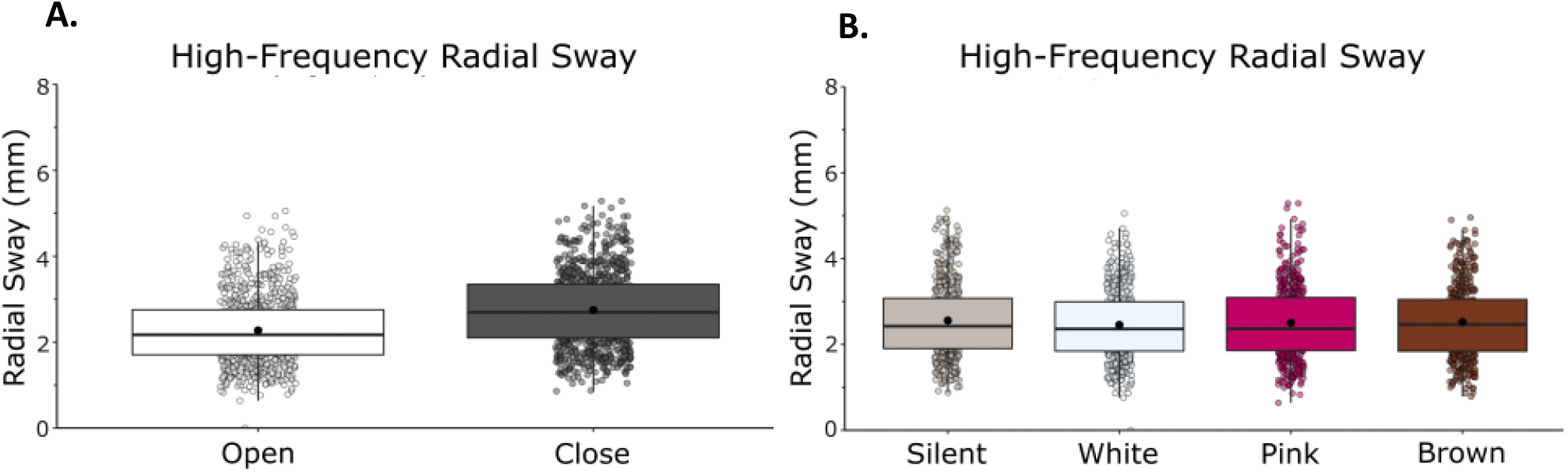
High-frequency (>0.3 Hz) RS was reduced with eyes open and there were no differences between white noise, pink noise or brown noise. A) High-frequency RS in eyes closed/eyes open. B) High-frequency RS in silent, white, pink, and brown noise conditions. Vision and noise contributed interactively to high-frequency RS. There was also an interaction effect between vision and stimulation. Box and whiskers plot with the solid black line representing the median, the solid black dot representing the mean, and the extending lines showing the maximum and minimum values.

Bonferroni-corrected post-hoc comparisons were performed to compare the individual noise condition effects on RS when compared to silence and to other noise conditions. When compared to silence (M = 2.56, SD = 0.87), there was no difference in RS when using white noise (M = 2.46, SD = 0.80, p = .083), pink noise (M = 2.50, SD = 0.85, p = .618), or brown noise (M = 2.52, SD = 0.79, p = 1.00). There was no difference between noise conditions when compared to each other: white and pink (p = 1.000), white and brown (p = .395), pink and brown (p = 1.000). The significant main effect of condition is not supported by the post-hoc tests.

### Low-Frequency Radial Sway

Low-frequency RS was reduced with vision and with noise (Fig. 3). We observed a main effect of vision (F(1,21) = 6.44, η = .23, p = .019) and a main effect of condition (F(1,21) = 2.79, η = .12, p = .048) on low-frequency RS (Fig. 3). We did not observe a vision by noise interaction (F(1,21) = 0.6867, η = .03, p = .563).

**Fig. 3.**
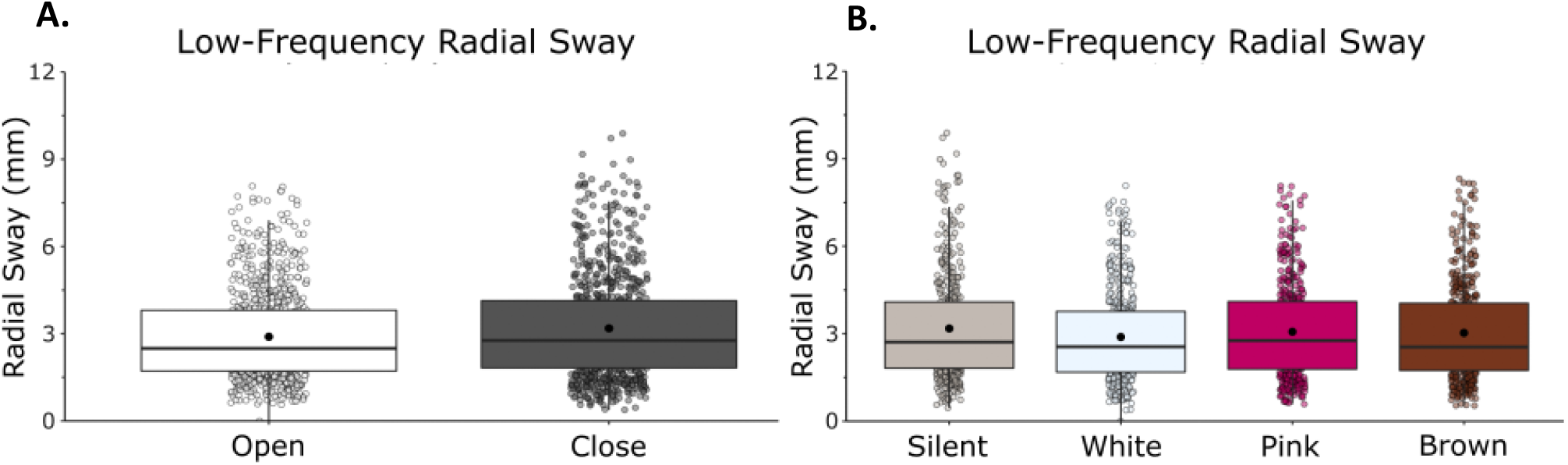
Low-frequency (<0.3 Hz) RS was reduced with eyes open and there were no differences between white noise, pink noise or brown noise. A) Low-frequency RS in eyes closed/eyes open. B) Low-frequency RS in silent, white, pink, and brown noise conditions. There was no interaction effect between vision and condition. Box and whiskers plot with the solid black line representing the median, the solid black dot representing the mean, and the extending lines showing the maximum and minimum values.

Bonferroni-corrected post-hoc comparisons were used to compare the individual conditions to each other. There was no difference between silence (M = 3.17, SD = 1.78) and white noise (M = 2.89, SD = 1.58, p = .208), pink noise (M = 3.07, SD = 1.66, p = 1.000) or brown noise (M = 3.03, SD = 1.68, p = 1.000). There was no difference between the stimulation conditions when compared to each other: white and pink (p = .548), white and brown (p = .993) and pink and brown (p = 1.000). The significant main effect of condition is not supported by the post-hoc tests.

### Detrended Fluctuation Analysis

Detrended Fluctuation Analysis showed that RS exhibits anti-persistent fractional Brownian motion (fαm, 1<α<1.5). Within this 1-1.5 range, we report differences between conditions in α. We observed a main effect of vision on α (F(1,21) = 39.11, η = .65, p = 0.001) (Fig. 4A) but no effect of condition on α (F(1,21) = 2.71, η = .11, p = 0.052) (Fig. 4B), indicating that sway patterns move in successive steps in random directions (semi-random walk) and tend toward the same direction to a higher degree during eyes open conditions than eyes closed conditions. We did not observe a vision by noise interaction (F(1,21) = 0.81, η = .04, p = .492).

**Fig. 4.**
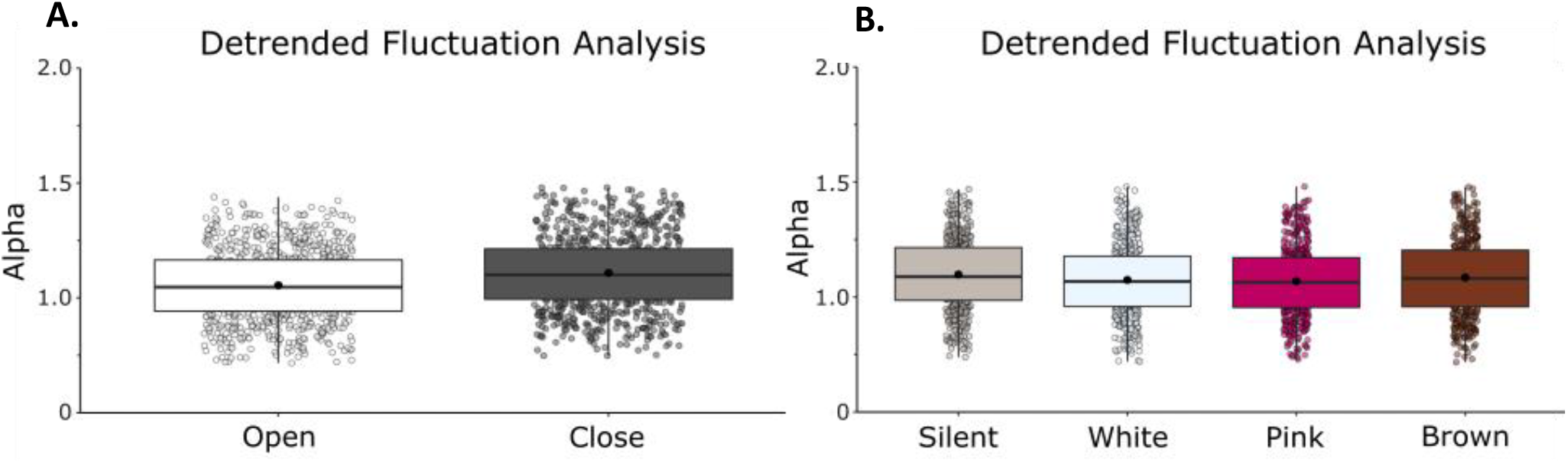
Detrended fluctuation analysis revealed a difference in the random-walk pattern commonly seen in postural sway between eyes open and eyes closed conditions. When eyes were closed there was an increase in alpha within the typical random-walk range. A) Mean α in eyes closed/eyes open B) Mean α in silent, white, pink, and brown noise conditions. Box and whiskers plot with the solid black line representing the median, the solid black dot representing the mean, and the extending lines showing the maximum and minimum values.

## Discussion

In Experiment 1, we did not find a convincing effect of the noise stimulation on postural sway variability. All stimuli were presented at 35 dB, which is above the noticeable threshold of the human ear but considered ‘quiet’ when considering that normal conversation happens at around 75 dB. We observed significant differences between the visual conditions, with eyes open causing a decrease when compared to eyes closed in radial sway variability, the high- and low-frequency components of radial sway, and DFA. However, the noise stimulation caused no changes to postural sway dynamics regardless of the noise stimuli structure. Even with the lack of an effect with auditory stimulation, this experiment further validated the importance of visual input on postural sway.

The lack of an effect that the stimulation condition had on postural sway may be explained by the optimization curve of SR. As previously mentioned, an optimal amount of added noise results in the maximal enhancement of behavioral performance, with further increases in intensity degrading the performance of that behavior (Moss, et al., 2004), and too small of an intensity of noise eliciting no information transfer or changes in behavior. At 35 dB, the noise may not be strong enough to elicit any behavioral changes during stimulation and may show that the changes seen in previous work on additive noise (Ross, et al., 2015, 2016; Priplata, et al., 2002, 2004, 2006) may be due to the SR phenomenon and not an attentional focus as commonly posited.

### Experiment 2

In Experiment 1 the intensity of noise stimulation may have been below the effective auditory threshold necessary to elicit the RS reduction reported in previous work with additive auditory noise (Ross et al. 2015, 2016; Carey et al. 2023). As previously discussed, the optimization curve of additive noise to the motor system may be relevant to effects on balance variability. Experiment 2 is a follow up study examining whether there are effects of vision and noise on RS, as in Experiment 1 but at a higher intensity of stimulation (75 dB).

## Results

### Radial Sway

We observed a main effect of vision (F(1,23) = 39.23, η = .63, p = .001) and a main effect of condition (F(1,23) = 11.94, η = .34, p = .001) on RS (Fig. 5). We did not observe a vision by noise interaction (F(1,23) = 1.12, η = .05, p = .346).

**Fig. 5.**
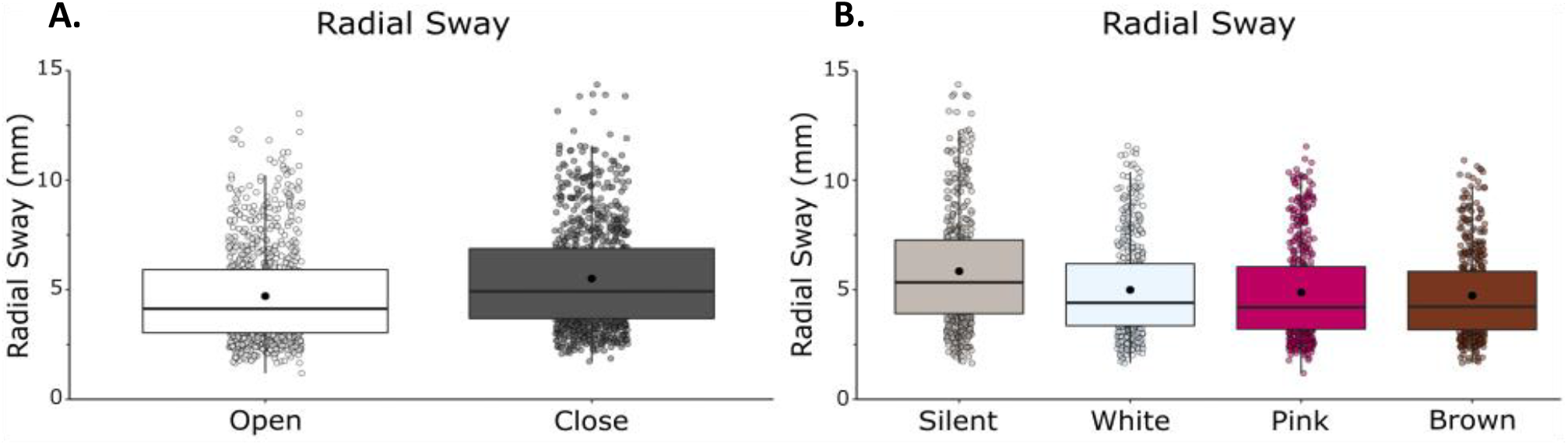
RS is significantly reduced with eyes open, with white noise, pink noise, and brown noise. A) RS in eyes closed/eyes open. B) RS in silent, white, pink, and brown noise conditions. There was no interaction effect between vision and condition. Box and whiskers plot with the solid black line representing the median, the solid black dot representing the mean, and the extending lines showing the maximum and minimum values.

Bonferroni-corrected post-hoc comparisons were performed to compare the individual stimulation conditions effects on RS when compared to silence and to other noise conditions. Post-hoc comparisons revealed a significant difference between silence (M = 5.84, SD = 2.58) and white noise (M = 4.98, SD = 2.13, p = .019), pink noise (M = 4.87, SD = 2.20, p = .011) and brown noise (M = 4.73, SD = 1.93, p = .002), confirming the main effect of condition in the ANOVA. We found no difference between stimulation conditions when compared to each other: white and pink (p = 1.00), white and brown (p = .149), pink and brown (p = .912).

### High-Frequency Radial Sway

We observed a main effect of vision (F(1,23) = 56.98, η = .71, p = .001) and a main effect of condition (F(1,23) = 11.66, η = .34, p = .001) on high-frequency RS (Fig. 6). We did not observe a vision by noise interaction (F(1,23) = 2.33, η = .09, p = .082).

**Fig. 6.**
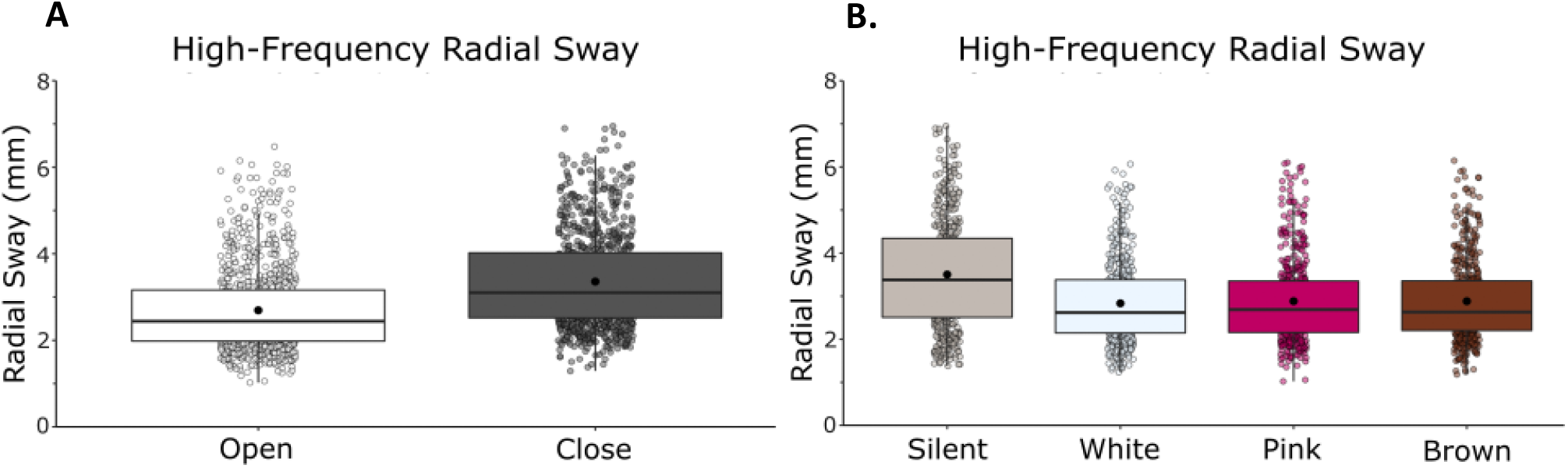
High-frequency (>0.3 Hz) sway was reduced with eyes open, with white noise, pink noise and brown noise. A) High-frequency RS in eyes closed/eyes open. B) High-frequency RS in silent, white, pink, and brown noise conditions. There was no interaction effect between vision and condition. Box and whiskers plot with the solid black line representing the median, the solid black dot representing the mean, and the extending lines showing the maximum and minimum values.

Bonferroni-corrected post-hoc comparisons were performed to compare the individual stimulation condition effects on RS when compared to silence and to other noise conditions. Post-hoc comparisons revealed a significant difference between silence (M = 3.50, SD = 1.22) and white noise (M = 2.83, SD = 0.92, p = .007), pink noise (M = 2.88, SD = 0.95, p = .018), and brown noise (M = 2.87, SD = 0.88, p = .009), confirming the main effect of condition in the ANOVA. There was no difference in effect between the noise conditions when compared to each other: white and pink (p = .568), white and brown (p = 1.000), pink and brown (p = 1.000).

### Low-frequency Radial Sway

We observed a main effect of vision (F(1,23) = 17.48, η = .43, p = .001) and a main effect of condition (F(1,23) = 8.86, η = .28, p = .001) on low-frequency RS (Fig. 7A and 7B). We did not observe a vision and noise interaction (F(1,23) = 0.66, η = .03, p = .576).

**Fig. 7.**
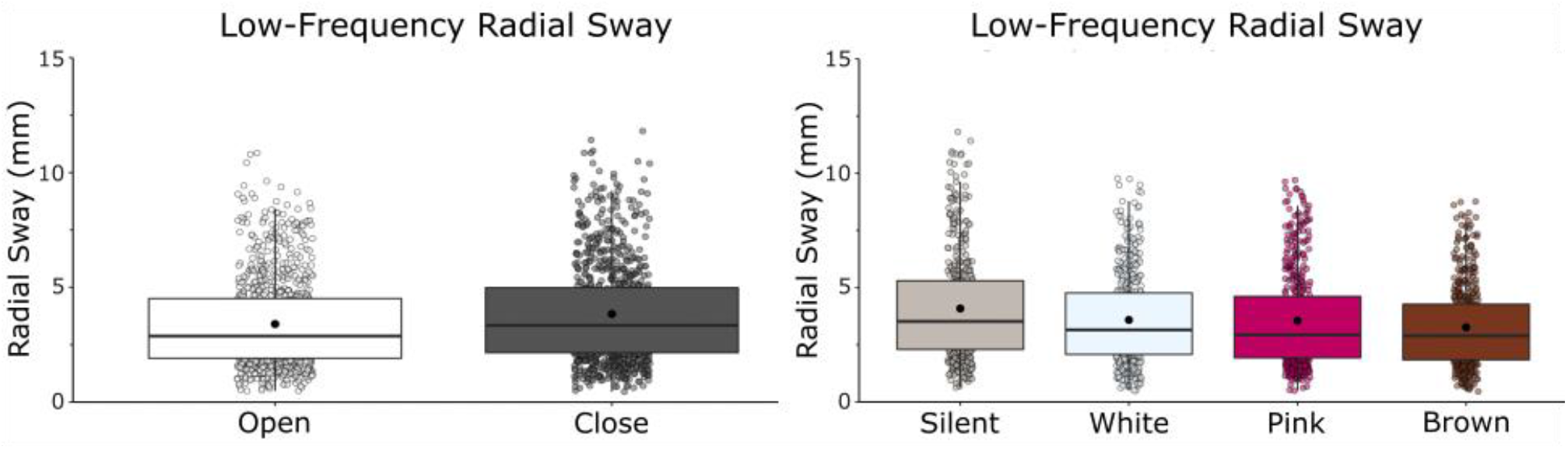
Low-frequency (<0.3 Hz) sway was reduced with eyes open and with brown noise. A) Low-frequency RS in eyes closed/eyes open. B) Low-frequency RS in silent, white, pink, and brown noise conditions. There was no interaction effect between vision and stimulation. Box and whiskers plot with the solid black line representing the median, the solid black dot representing the mean, and the extending lines showing the maximum and minimum values.

Bonferroni-corrected post-hoc comparisons were performed to compare the individual stimulation condition effects on RS when compared to silence and to other noise conditions. Post-hoc comparisons revealed a significant difference between silence (M = 4.08, SD = 2.33) and brown noise (M = 3.26, SD = 1.77, p = .016), but no effect of white (M = 3.58, SD = 1.93, p = .073) or pink noise (M = 3.54, SD = 2.11, p = .092). There was no difference in effect between the noise conditions when compared to each other: the white and pink (p = 1.000), white and brown (p = .071) and pink and brown (p = .124).

### Detrended Fluctuation Analysis

Detrended Fluctuation Analysis showed that our RS data exhibit anti-persistent fractional Brownian motion (fαm, 1<α<1.5). We observed a main effect of vision on α (F(1,21) = 13.94, η?= .38, p = 0.001) (Fig. 8A) and a main effect of noise on α (F(1,21) = 3.97, η = .15, p = 0.01) (Fig. 4B), indicating that with visual input and with noise sway patterns tend toward a more positive correlation than without visual input or noise. We did not observe a vision by noise interaction (F(1,21) = 0.16, η = .01, p = .923).

**Fig. 8.**
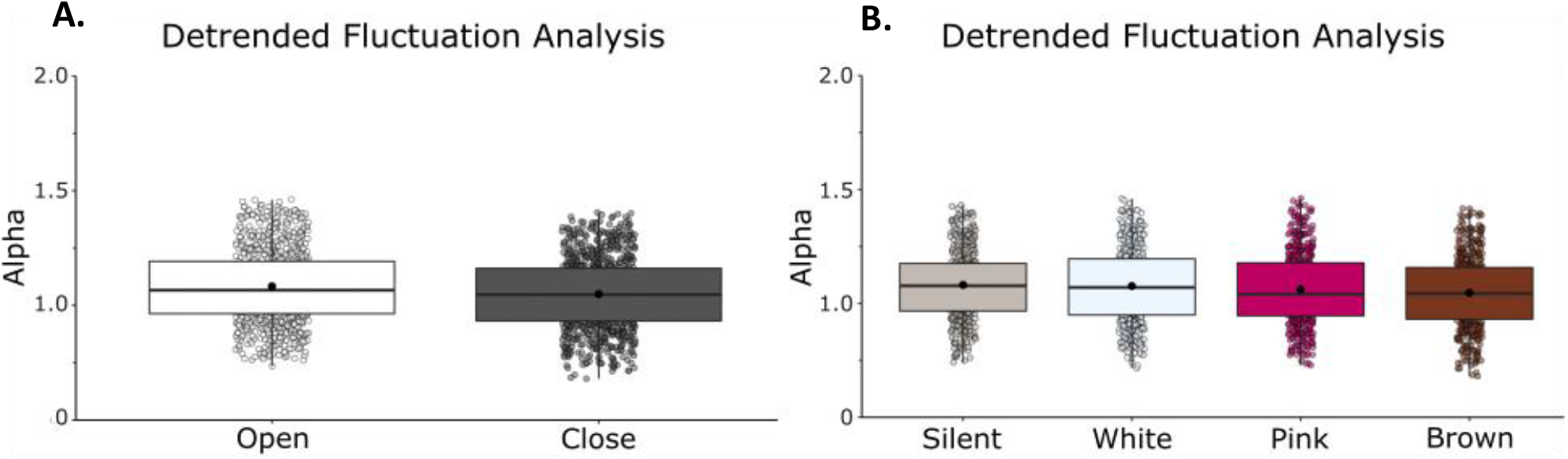
Detrended fluctuation analysis revealed changes with vision and brown noise stimulation in the random-walk pattern of sway. A) Mean α in eyes closed/eyes open B)

Bonferroni-corrected post-hoc comparisons were performed to compare the individual stimulation condition effects on RS when compared to silence and to other noise conditions. Post-hoc comparisons revealed a significant difference between silence (M = 1.081, SD = 0.149) and brown noise (M = 1.047, SD = 0.158, p = 0.027), but not white (M = 1.076, SD = 0.164, p = 1.000) or pink noise (M = 1.061, SD = 0.161, p = .5644). This finding suggests that there is a shift in the correlational structure of sway specific to brown noise stimulation. During brown noise, the scaling exponent becomes closer to α = 1.0 indicating a more negative correlation of the signal, and therefore a more random path of movement. When comparing the stimulation conditions to each other, we found a significant difference between white and brown noise (p = 0.027), but no differences between white and pink (p = 1.000) or pink and brown (p = 1.000).

Mean α in silent, white, pink, and brown noise conditions. Box and whiskers plot with the solid black line representing the median, the solid black dot representing the mean, and the extending lines showing the maximum and minimum values.

## Discussion

Experiment 2 showed the sway reducing influence that auditory noise stimulation can have on the postural system. Presented at 75 dB, all three noise stimulation conditions caused reductions in the averaged radial sway and the high- and low-frequency components of radial sway. The results from Experiment 2 support that postural sway variability is decreased with visual input and with auditory stimulation.

More interestingly, the DFA scaling exponent was impacted by the structure of the noise presented. Stability can be understood as the co-adjustment of local variability and serial correlational properties (Blázquez et al., 2010). In this study, DFA of sway from participants revealed a lower scaling coefficient (α) when brown noise was introduced and when visual input was available. Higher α indicates more persistence, or more correlation between successive points of sway, and a lower α indicates more anti-persistence in the sway. Anti-persistence can be interpreted as more tightly controlled, or less resistant to changes in COP displacement direction, which reflects adaptability of the signal to change (Ducharme and van Emmerik, 2018).

Our results contribute to the knowledge about variability and adaptability by suggesting that the reduction in sway variability with brown noise specifically is accompanied by a potential increase in the adaptability of the postural system. Importantly, however, we emphasize that α was between 1 and 1.5 in all noise stimulation conditions; sway remained anti-persistent and the differences between conditions only show differences between the degree of anti-persistence within this range. Auditory noise stimulation did not interfere with the random walk property of sway, but it may have influenced adaptability as well as variability leading to decreased postural sway.

The reductions seen in radial sway during auditory noise can be at least partially explained by the SR phenomenon. The theory of SR explains the amplification of information-carrying signals through the addition of noise (Hängii, 2002). This phenomenon relies on a threshold-based system (the postural system), additive noise (auditory stimuli), and an underlying information carrying signal (motor responses of the postural system), all of which were present within this study. However, other possible explanations for the noise effect on postural sway is that there is an increased attentional arousal during stimulation conditions. Our results also suggest that there is frequency matching that could have a functional impact on balance variability and adaptability. More work is needed to determine which mechanisms contribute most to this shift in behavioral output during noise stimulation.

## General Discussion

In both experiments, postural variability was significantly reduced with visual input, helping to validate the importance of visual sensory information on the postural control system. However, there was a significant difference in the effect of noise stimulation between the two experiments. During Experiment 1, there was no effect of the noise stimulation on postural sway variability. Although our ANOVAs showed main effects of condition, pairwise comparisons showed no effect between noise and silence. The stimulation was presented at 35 dB, which is above the noticeable threshold of human hearing but is considered ‘quiet’ compared with intensities of around 75 dB, which is slightly higher than human speech (*What Noises Cause Hearing Loss?* | *NCEH* | *CDC*., 2022, November 8). Experiment 2 showed the potential impact that auditory noise can have on our postural sway system. Presented at 75 dB, instead of 35 dB as in Experiment 1, the noise stimulation caused a reduction in radial sway when compared to silence, as well as when it was separated into high- and low-frequencies (Table 1 for effect sizes from both experiments). We found no significant differences in RS between the three noise conditions in either experiment. The results from Experiment 2 support that postural sway variability is decreased with visual input and with noise stimulation, regardless of the structure of the noise signal.

**Table 1.**
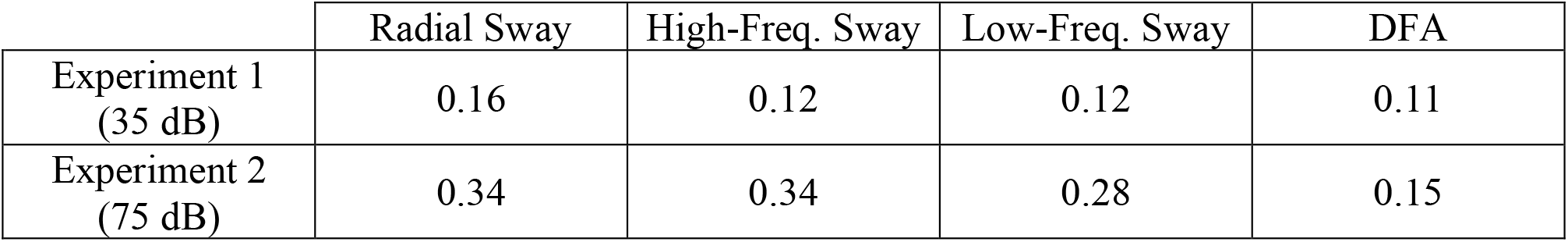
Effect sizes of the comparisons across noise conditions from the ANOVAs of each experiment. An effect size (f) of > 0.4 was considered to reflect a strong effect (Cohen, 1988).

|  | Radial Sway | High-Freq. Sway | Low-Freq. Sway | DFA |
| --- | --- | --- | --- | --- |
| Experiment 1<br>(35 dB) | 0.16 | 0.12 | 0.12 | 0.11 |
| Experiment 2<br>(75 dB) | 0.34 | 0.34 | 0.28 | 0.15 |

When observing RS of the high- and low-frequencies of sway, we found that there are independent impacts of the different noise structures on sway. Work by van den Heuvel *et al*. (2009) and further established by Yeh *et al*. (2010) showed that sensory feedback can affect these low and high frequency components of sway differentially. The slower timescales of sway, reflecting drift of the inertial mass of the body (Winter et al., 1998), are more susceptible to abrupt changes in sensory feedback (Yeh et al., 2010, 2014; van den Heuvel et al., 2009). While faster timescales of sway, reflecting small adjustments of the center of mass used to maintain stability, are susceptible to the joint rigidity and muscle activations (Kiemel et al., 2005; Peterka, 2002). By separating the components of postural sway into low- and high-frequencies we are able to examine the dynamics of sway more thoroughly to discern if specific dynamics are more heavily influenced by the structure of the noise signals presented. Our results of Experiment 1 support that there was no influence of noise on the separate frequency components of postural sway. However, in Experiment 2 we discovered that additive noise can decrease the radial sway variability in both the low- and high-frequencies. There were no differences between the white, pink, or brown noise in these frequency components, but all three noise signals reduced RS when compared to silence.

The differences in results between Experiment 1 and Experiment 2 were unexpected. However, if we interpret the results while considering the theory of stochastic resonance this effect of intensity is precedented. Past work on SR has shown the influence of the intensity of additive noise for altering the behavioral output of the system under study (Moss et al, 2004; van der Groen, 2018). As previously explained, within the theory of SR added noise helps to enhance the underlying information signal of interest. However, too much added noise to the information signal of interest can cause the signal to become hidden. An optimal amount of added noise results in the maximum enhancement of behavioral output, whereas further increases in the noise intensity only degrade the detectability or information content. Similarly, noise at too low of an intensity may elicit no changes in performance (Moss, et al. 2004). This relationship between intensity of noise and SR is not linear. As for the interest of this study, Experiment 1 and 2 differed in only one way: the intensity at which the noise stimuli were presented. Experiment 1 presented stimuli at 35 dB while Experiment 2 presented the stimuli at 75 dB. This difference in noise intensity may explain the differences we see in the results. Too low of a noise intensity may fail to result in alterations in the behavioral output of the system being stimulated. Using a higher intensity elicits the behavioral output expected: a reduction in radial sway variability and change in the dynamics of postural sway.

Another possible explanation for the noise effect on postural sway is that there is an increased attentional arousal during auditory stimulation, which could lead to a higher level of control in sway. Cluff et al. (2010) showed that adding a cognitive task during quiet standing leads to an increase in the automaticity of the postural system and to improvements of stability. However, it has also been shown that passively listening to a single sustained auditory tone does not affect postural sway (Deviterne et al., 2005), so we would not predict that auditory attention in our sustained noise conditions would drive a stabilizing effect in the current experiment. Similarly, if an attentional mechanism was causing this effect, we would expect to see a reduction in RS in both experiments, not just the one of higher intensity. However, it could be the case that lower intensity noise has less attentional demands during the task.

Although the theory of SR at least partially explains our current results, more research is required to determine the specific mechanisms driving this reduction in sway variability. Whether or not these effects are due to SR, attention, or some other mechanism, the findings have profound implications for improving balance in populations at high-risk for falls. One explanation for the minimal effects of structure (white, pink, brown) on the noise effect is that our sensory systems processes and utilize all forms of ‘noise’ in the same way. In our daily environment, humans are exposed to multiple different types of noise of varying structures and intensities. From the wind blowing through trees to the sound of cars on a road, naturally occurring noise can vary in many ways, but our postural or sensory systems may have become insensitive to these variations of frequency content for functional incorporation into the reduction of movement variability. However, the DFA results specific to brown noise in Experiment 2 suggest this not to be the case for all aspects of movement.

## Limitations

The introduction of auditory noise in our experimental setting may be challenging to translate into a clinical or home setting. However, there is a multitude of sources of noise within our environment that may be influencing our motor systems without our conscious awareness. Additionally, due to this paper covering two separate experiments performed at different times, the comparison of the two studies is not as straightforward. Future work should assess the intensity of auditory noise stimulation within a single experimental paradigm to elucidate the influence intensity of stimulation may have on our motor system.

## STAR ★ METHODS

### Resource availability

#### Lead contact

Questions and requests for information and data/code should be directed to the corresponding author.

#### Data and Code Availability

- All data reported in this paper will be shared by the lead contact upon request.
- Code used for stimulus presentation and code used to analyze the data can be requested from the *lead contact*.
- Any additional information needed to assess the current behavioral data can be obtained via contacting the *lead contact*.

## Experimental Model and Subject Details

### Methods

#### Experimental Design

The current study included two within-subject experiments that were conducted three months apart. No subject participated in both experiments. The intention was to study how auditory stimulation and the structure of noise signals may influence postural control during standing. First, we tested 3 noise types using a low intensity of 35 dB. We then ran the second experiment at a higher intensity in an attempt to understand whether the intensity of noise amplitude may have influenced the results of Experiment 1. A power analysis with a strong effect size (> 0.4) when using a repeated measured ANOVA with two levels (eyes open vs. eyes closed) and four conditions (silence vs. white vs. pink vs. brown) was performed for each study that resulted in an approximate 25 participants needed to observe a significant effect size.

#### Participants in Experiment 1

Twenty-four healthy young adults, 7 male and 17 female, (mean age = 20.61 ± 2.87 years) of varying heights (64.63 ± 5.15 inches) and weights (148.71 ± 25.85 lbs.) were recruited from the University of California, Merced student population. Self-report screeners were used to exclude participants with hearing impairments, arthritis, orthopedic conditions, or neurological disorders (Ross *et al*. 2015, 2016). No participants reported recent injuries or skeletomuscular disorders, and all could stand unassisted during the experiment. The experimental protocol was carried out in accordance with the Declaration of Helsinki, reviewed by the UC Merced IRB, and all participants gave their informed and written consent prior to testing.

Participants were instructed to stand on a force platform in a relaxed, comfortable standing position with their arms at their sides while wearing headphones through which the auditory stimuli were presented. Participants were instructed to keep their eyes fixated on a black crosshair stimulus posted on the wall 229 cm in front of them at approximately eye level for the eyes-open trials and to keep their head facing forward and eyes closed during eyes closed trials ( Ross et al., 2015, Ross et al., 2016).

The noise (silence, white, pink, brown) and visual input (eyes open, eyes closed) conditions were presented in a randomized order. There were a total of 80 trials, 20 trials for each noise condition, 10 with eyes opened and 10 with eyes closed. The trials lasted 20 seconds and were accompanied by silence, auditory white noise, auditory pink noise, or auditory brown noise. The noise was presented at an intensity of 35 dB in Experiment 1 and 75 dB in Experiment 2. Center of Pressure (CoP) was sampled at 200 Hz with an AMTI Force and Motion platform (Optima BP400600-2000 GEN 5; AMTI Force and Motion, Watertown, MA, USA). All data for each subject were collected in a single session. The auditory noise stimuli were generated using MATLAB to be random signals with a constant spectral density. Participants were exposed to the noise stimuli prior to the experiment to verify that the intensity was not uncomfortable for them. No participants reported discomfort at these intensities.

#### Participants in Experiment 2

Twenty-two healthy young adults, 9 male and 13 female, (mean age = 21.96 ± 3.42 years) of varying heights (65.56 ± 3.48 inches) and weights (141.76 ± 27.28 lbs.) were recruited from the University of California, Merced student population. Self-report screenings were used to exclude participants with hearing impairments, arthritis, orthopedic conditions, or neurological disorders (following Ross et al., 2015, 2016). No participants reported recent injuries or skeletomuscular disorders, and all could stand unassisted during the experiment. The experimental protocol was carried out in accordance with the Declaration of Helsinki, reviewed by the UC Merced IRB, and all participants gave their informed and written consent prior to testing.

### Analysis

The CoP of each condition was analyzed using custom scripts in MATLAB (MathWorks, Natick, MA, USA). The first 4 seconds of each trial were removed to eliminate any potential startle response that participants might have had to stimulus onset. Radial sway (RS) of the CoP was calculated for each sample (i) using the anterior-posterior (A-P; x) and medial-lateral (M-L; y) components of sway following (Lafond et al., 2004a, b):

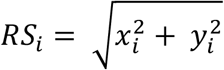

Average RS was calculated for each trial and was used to assess bidirectional variability in CoP during the trials (Lafond et al., 2004a, b). Although RS is not a direct metric of stability, it utilizes the multidirectional variability of sway to offer a more robust understanding of the sway dynamics which may lead to stability (Lafond et al., 2004a, b). Trial outliers were determined as trials with averages of ±2 standard deviations from that subject’s mean within condition and were removed. In Experiment 1, 203 of the total 5280 trials were removed, resulting in a removal rate of 4%. In Experiment 2, 295 of the total 5,760 trials were removed resulting in a 5% removal rate. Participants were excluded if they were suspected of making movements other than postural sway during data collection (Edwards, 1946; Hafström et al., 2002).

In each Experiment, the effects of noise during eyes opened and eyes closed on mean RS were modeled across conditions using a two (eyes open vs. eyes closed) × four (silence vs. white vs. pink vs. brown) analysis of variance with repeated measures and with subjects as a between factor.

The analysis was then repeated using the filtered high and low frequency RS separately to assess changes in slower and faster timescales of postural control (following the methods of Yeh et al., 2010 and Yeh et al., 2014). Postural sway is naturally oscillatory and is composed of two primary timescales of oscillation (Yeh et al., 2010). Low-frequency oscillations are typically considered to reflect feedback-based corrective responses, where high-frequency oscillations are considered open-loop exploratory processes (Yeh et al., 2014). We used low- and high-pass Butterworth filtering routines, as in Yeh et al., 2014, to decompose sway into low (<0.3 Hz) and high (>0.3 Hz) frequency sway. The filter cutoff was chosen based on van den Heuvel et al., 2009 and Jeka, et al., 1997 to separate into sensory feedback-related sway and spontaneous/exploratory sway.

Detrended fluctuation analysis (DFA) was used to assess the sway dynamics over time while under different stimulation conditions (Deligniéres, et al., 2002; Collins et al., 1994). DFA is used to study the behavior of the timeseries of CoP. This analysis, first introduced by Peng et al., (1994), is a scaling analysis method that provides a scaling exponent α, which offers information about the correlational properties of the CoP signal. When the DFA value exists between 1 < α <1.5, the postural sway is considered antipersistent. This means that the sway moves in successive steps in random directions (a semi-random walk) and does not trend toward the same direction. The scaling exponent α includes the information concerning the correlation properties of the signal: α = 1.5 is characteristic of an uncorrelated random series (white noise), while the signal presents positive correlations if α > 1.5 and negative correlations if α < 1.5. Antipersistent radial sway dynamics are commonly described in healthy postural sway. This analysis was completed as in (Blázquez et al., 2003) using the same parameters. See Blázquez et al., 2010 and Deligniéres et al., 2003 for more details on the DFA method.

## Author contributions

Conceptualization, J.M.R and D.A.; Methodology, A.J.S, K.C.B., H.B., and M.G.G.; Investigation, S.C.; Formal Analysis, S.C..; Writing – Original Draft, S.C. Writing – Review & Editing, S.C., J.M.R., D.A., and R.B..; Visualization, S.C.: Supervision, R.B.

## Declaration of Interest

The authors declare no competing interests.

## Notes

### Competing Interest Statement

The authors have declared no competing interest.

